# Bulk Tissue Cell Type Deconvolution with Multi-Subject Single-Cell Expression Reference

**DOI:** 10.1101/354944

**Authors:** Xuran Wang, Jihwan Park, Katalin Susztak, Nancy R. Zhang, Mingyao Li

**Affiliations:** Graduate Group in Applied Mathematics and Computational Science, University of Pennsylvania, Philadelphia, PA; Departments of Medicine and Genetics, University of Pennsylvania, Philadelphia, PA; Department of Statistics, The Wharton School, University of Pennsylvania, Philadelphia, PA; Department of Biostatistics, Epidemiology & Informatics, University of Pennsylvania, Philadelphia, PA

**Author notes:** Correspondence: Nancy R. Zhang Mingyao Li.

## Abstract

We present MuSiC, a method that utilizes cell-type specific gene expression from single-cell RNA sequencing (RNA-seq) data to characterize cell type compositions from bulk RNA-seq data in complex tissues. When applied to pancreatic islet and whole kidney expression data in human, mouse, and rats, MuSiC outperformed existing methods, especially for tissues with closely related cell types. MuSiC enables characterization of cellular heterogeneity of complex tissues for identification of disease mechanisms.

---

Bulk tissue RNA-seq is a widely adopted method to understand genome-wide transcriptomic variations in different conditions such as disease states. Bulk RNA-seq measures the average expression of genes, which is the sum of cell type-specific gene expression weighted by cell type proportions. Knowledge of cell type composition and their proportions in intact tissues is important, because certain cell types are more vulnerable for disease than others. Characterizing the variation of cell type composition across subjects can identify cellular targets of disease, and adjusting for these variations can clarify downstream analysis.

The rapid development of single-cell RNA-seq (scRNA-seq) technologies have enabled cell type-specific transcriptome profiling. Although cell type composition and proportions are obtainable from scRNA-seq, scRNA-seq is still costly, prohibiting its application in clinical studies that involve a large number of subjects. Furthermore, scRNA-seq is not well suited to characterizing cell type proportions in a solid tissue, because the cell dissociation step is biased towards certain cell types^1^.

Computational methods have been developed to deconvolve cell type proportions using cell type-specific gene expression references^2^. CIBERSORT^3^, based on support vector regression, is a widely used method designed for microarray data. More recently, BSEQ-sc^4^ extended CIBERSORT to allow the use of scRNA-seq gene expression as a reference. TIMER^5^, developed for cancer data, focuses on the quantification of immune cell infiltration. These methods rely on pre-selected cell type-specific marker genes, and thus are sensitive to the choice of significance threshold. More importantly, these methods ignore cross-subject heterogeneity in cell type-specific gene expression as well as within-cell type stochasticity of single-cell gene expression, both of which cannot be ignored based on our analysis of multiple scRNA-seq datasets (**Supplementary Figure 1a**).

Here we introduce a new <u>MU</u>lti-Subject <u>Si</u>ngle <u>C</u>ell deconvolution (MuSiC) method (https://github.com/xuranw/MuSiC) that utilizes cross-subject scRNA-seq to estimate cell type proportions in bulk RNA-seq data (**Figure 1**). A key concept in MuSiC is “marker gene stability”. We show that, when using scRNA-seq data as a reference for cell type deconvolution, two fundamental types of stability must be considered: cross-subject and cross-cell, in which the first is to guard against bias in subject selection, and the second is to guard against bias in cell capture in scRNA-seq. By incorporating both types of stability, MuSiC allows for scRNA-seq datasets to serve as effective references for independent bulk RNA-seq datasets involving different individuals.

**Figure 1:**
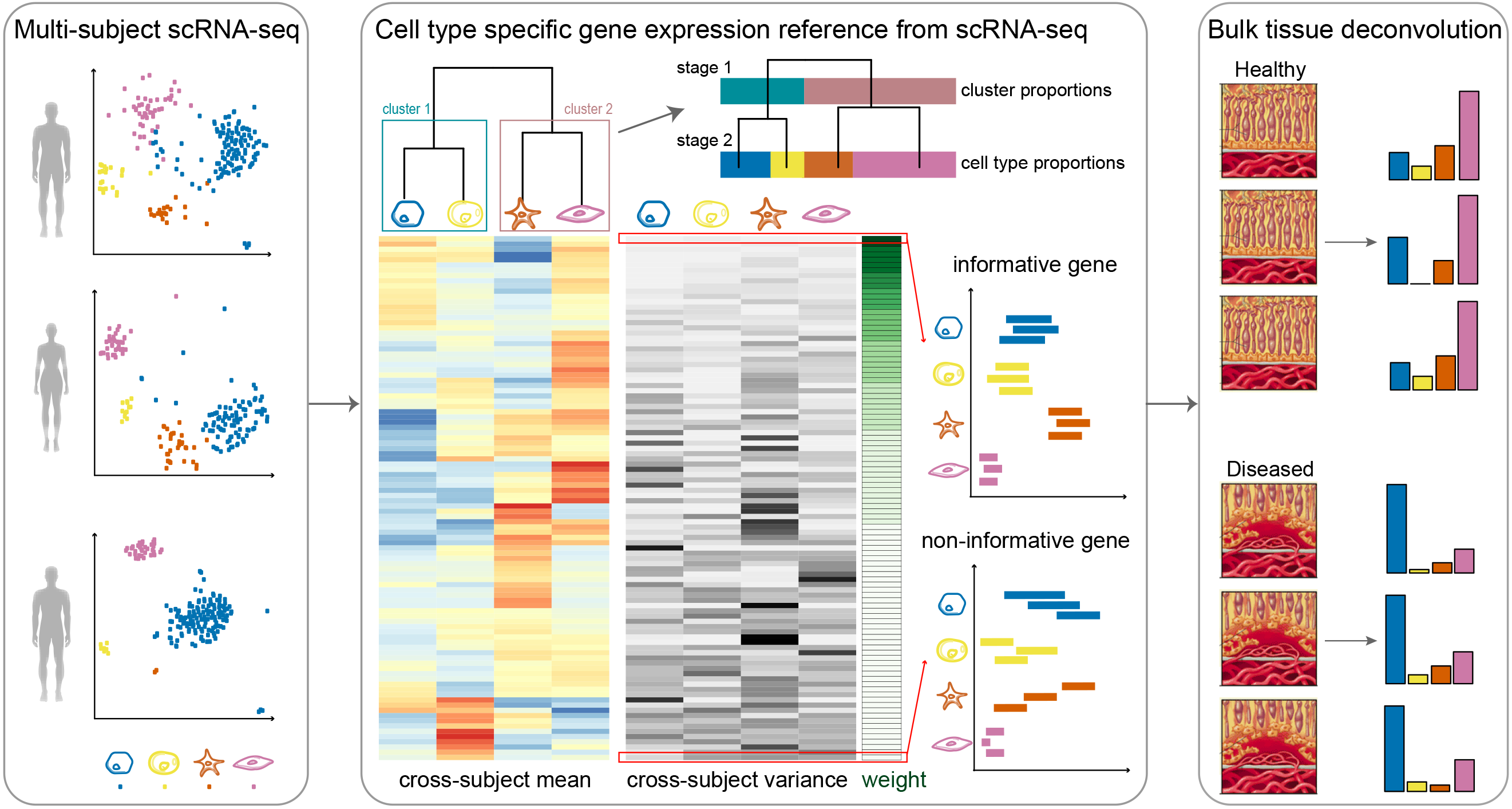
Overview of MuSiC framework. MuSiC starts from scRNA-seq data from multiple subjects, classified into cell types (shown in different colors), and constructs a hierarchical clustering tree reflecting the similarity between cell types. Based on this tree, the user can determine the stages of recursive estimation and which cell types to group together at each stage. MuSiC then determines the group-stable genes and calculates cross-subject mean (red to blue) and cross-subject variance (black to white) for these genes in each cell type. MuSiC up-weighs genes with low cross-subject variance and down-weighs genes with high crosssubject variance. In the example shown, deconvolution is performed in two stages, only cluster proportions are estimated for the first stage. Constrained by these cluster proportions, the second stage estimates cell type proportions, illustrated by the length of the bar with different colors. The deconvolved cell type proportions can then be compared across disease cohorts.

Rather than pre-selecting marker genes from scRNA-seq based only on mean expression, MuSiC gives weight to each gene, allowing for the use of a larger set of genes in deconvolution. The weighting scheme prioritizes stable genes across subjects: up-weighing genes with low cross-subject variance (informative genes) and down-weighing genes with high cross-subject variance (non-informative genes). This requirement on cross-subject stability is critical for transferring cell type-specific gene expression information from one dataset to another.

Solid tissues often contain closely related cell types, and correlation of gene expression between these cell types leads to collinearity, making it difficult to resolve their relative proportions in bulk data. To deal with collinearity, MuSiC employs a tree-guided procedure that recursively zooms in on closely related cell types. Briefly, we first group similar cell types into the same cluster and estimate cluster proportions, then recursively repeat this procedure within each cluster (**Figure 1**). At each recursion stage, we only use genes that have low within-cluster variance, a.k.a. the cross-cell stable genes. This is critical as the mean expression estimates of genes with high variance are affected by the pervasive bias in cell capture of scRNA-seq experiments, and thus cannot serve as reliable reference. See online methods for details.

To demonstrate and evaluate MuSiC, we started with a well-studied tissue, the islets of Langerhans, which are clusters of endocrine cells within the pancreas that are essential for blood glucose homeostasis. Pancreatic islets contain five endocrine cell types (α,β,δ,ϵ, and γ), of which β cells, which secrete insulin, are gradually lost during type 2 diabetes (T2D). We applied MuSiC to bulk pancreatic islet RNA-seq samples from 89 donors from Fadista et al.^6^, to estimate cell type proportions and to characterize their associations with hemoglobin A1c (HbA1c) level, an important biomarker for T2D. We were motivated to re-analyze this data because, as shown in **Figure 2** and in Baron et al.^4^, existing methods failed to recover the correct β cell proportions, which should be around 50-60%^7^, and also failed to recover their expected negative relationship with HbA1c level. As reference, we experimented with scRNA-seq data from two sources: 6 healthy and 4 T2D adult donors from Segerstolpe et al.^8^, and 12 healthy and 6 T2D adult donors from Xin et al.^9^. All bulk and single-cell datasets in this analysis are summarized in **Supplementary Table 1**.

**Figure 2:**
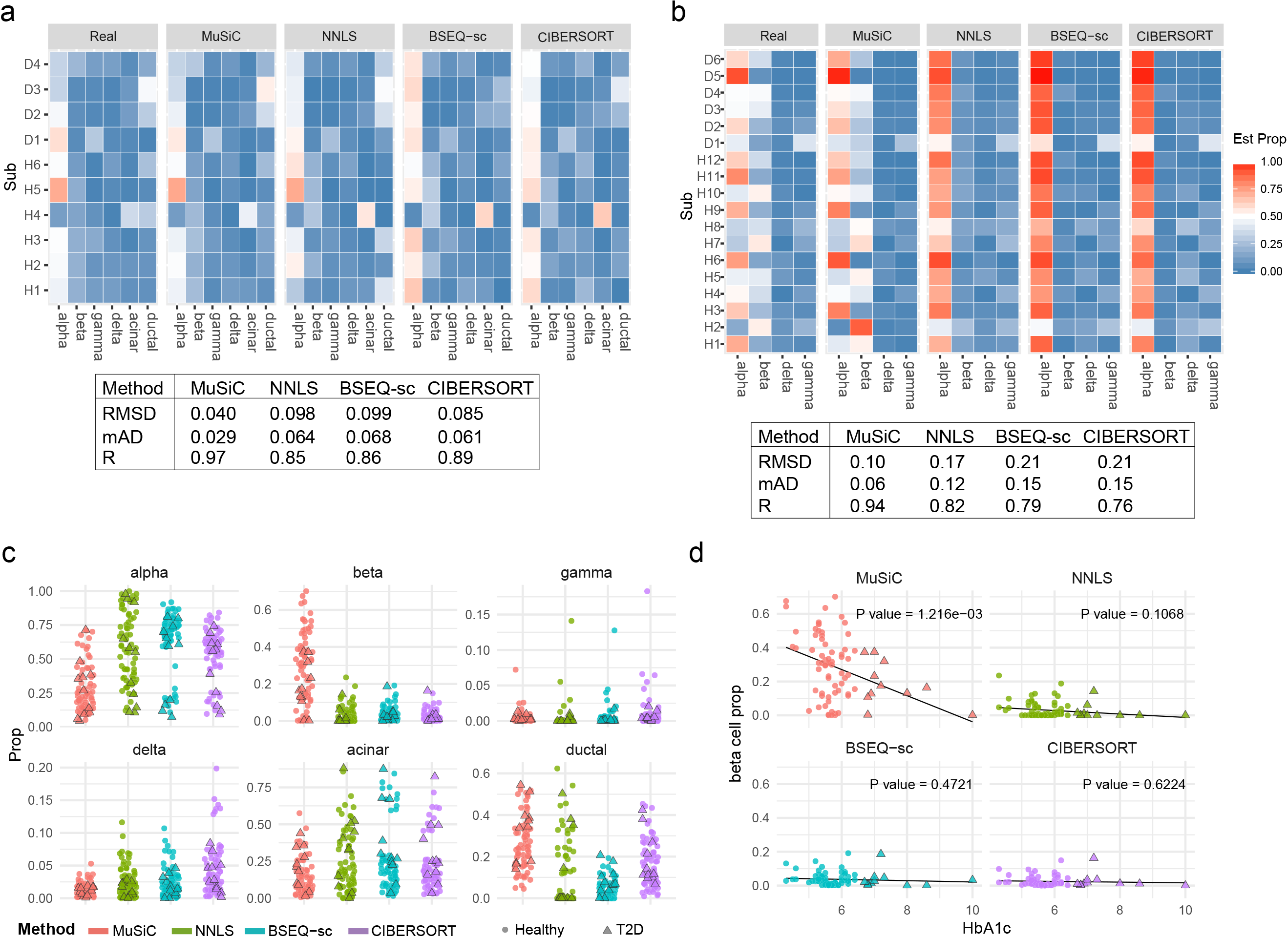
Pancreatic islet cell type composition in healthy and T2D human samples. **a** and **b** Benchmarking of deconvolution accuracy on bulk data constructed by combining together scRNA-seq samples. **a**. The bulk data is constructed for 10 subjects from Segerstolpe et al. while the single cell reference is taken from the same dataset. The cell type proportions of healthy subjects are estimated by leave-one-out single cell reference. The subject names are relabeled; the table shows average root mean square error (RMSD), mean absolute deviation (mAD), and Pearson correlation (R) across all samples and cell types. **b**. The bulk data is constructed for 18 subjects from Xin et al. while the single cell reference is 6 healthy subjects from Segerstolpe et al. **c**. Jitter plots of estimated cell type proportions for Fadista et al subjects, color-coded by deconvolution method. Of the 89 subjects from Fadista et al., only the 77 that have recorded HbA1c level are plotted, and T2D subjects are denoted as triangles. **d**. HbA1c vs beta cell type proportions estimated by each of 4 methods. The reported p-values are from single variable regression β cell proportion ~ HbA1c. Multivariable regression results are reported in **Supplementary Table 3**.

First, to systematically benchmark, we applied MuSiC and three other methods (Nonnegative least squares (NNLS), CIBERSORT, and BSEQ-sc) to artificial bulk RNA-seq data constructed by simply summing the scRNA-seq read counts across cells for each single-cell sequenced subject. In this case, true cell type proportions are known, which allows the evaluation of accuracy. More details on artificial bulk construction are described in the **Supplementary Note. Figure 2a, Supplementary Figure 1c** and **Supplementary Figure 2b** show the estimation results when the artificial bulk and the single-cell reference data are from the same study, either both from Segerstolpe et al.^8^ or both from Xin et al.^9^. MuSiC achieves improved accuracy over existing procedures. **Figure 2b** and **Supplementary Figure 2a** show the estimation results when the artificial bulk and the single-cell reference data are from different studies. This is a more challenging but more realistic scenario, since library preparation protocols vary across labs and bulk deconvolution analyses are often performed using single-cell reference generated by others. MuSiC still maintains high accuracy, while other methods perform substantially worse. Further comparisons show that, unlike existing methods that rely on pre-selected marker genes, MuSiC gives accurate results when the cell type composition in the bulk data is substantially different from that of the single cell reference (**Supplementary Figure 2c** and **Supplementary Note 2**), and when the bulk tissue contains minority cell types that are missing in the reference (**Supplementary Figure 3** and **Supplementary Note 3**). MuSiC’s ability to transfer knowledge across data sources is derived from its consideration of marker gene stability.

We now turn to the deconvolution of bulk RNA-seq data from Fadista et al.^6^. We used the scRNA-seq data from Segerstolpe et al. as reference for all methods. MuSiC recovers the expected ~50-60% β cell proportion for the healthy subjects^7^, whereas other methods grossly overestimate the proportion of α cells and underestimate the proportion of β cells. Furthermore, MuSiC detects a significant association of β cell proportion with HbA1c level (p-value 0.00126, **Figure 2d**). Based on clinical standard, HbA1c level <6.0% is classified as normal, and >6.5% is classified as diabetic. After adjusting for age, gender and body mass index, MuSiC estimates suggest that 0.5% increase in HbA1c level, representing the magnitude of increase from normal to the diabetes cutoff, corresponds to a drop of 6.14% ± 4.98% in β cell proportion.

As a second tissue example, we used the kidney, a complex organ consisting of several anatomically distinct segments each playing critical roles in the filtration and reabsorption of electrolytes and small molecules of the blood. Chronic kidney disease (CKD), the gradual loss of kidney function, is increasingly recognized as a major health problem, affecting 10-16% of the global adult population. We aim to characterize how kidney cell type composition changes during CKD. Fibrosis is the histologic hallmark common to all CKD models, and hence, we analyzed the bulk RNA-seq data from three mouse models for renal fibrosis: unilateral ureteric obstruction induced by surgical ligation of the ureter (UUO, Arvaniti et al.^10^), toxic precipitation in the tubules induced by high dose folic acid injection (FA, Craciun et al.^11^), or genetic alteration by transgenic expression of genetic risk variant APOL1 in podocytes (APOL1 transgenic mice^12^). As reference, we used the mouse kidney specific scRNA-seq data from Park et al.^1^. Details of all datasets are summarized in **Supplementary Table 2**. We systematically benchmarked all methods on artificial bulk experiments performed using the Park et al. scRNA-seq data, finding similar trends as those in **Figure 2a-b** (**Supplementary Figure 4a-b**).

Hierarchical clustering of the cell types in the single cell reference reveals that, apart from neutrophils and podocytes, kidney cells fall into two large groups: Immune cell types (macrophages, fibroblasts, T lymphocytes, B lymphocytes, and natural killer cells) and kidney-specific cell types (proximal tubule, distal convolved tubule, loop of Henle, two cell types forming the collecting ducts, and endothelial cells). Of these, proximal tubule (PT) is the dominant cell type in kidney, and the proportion of PT cells is known to decrease with CKD progression. MuSiC finds this decrease in all three mouse models (**Figure 3b-d**). Other methods also detect this association for the APOL1 and UUO mouse models, but showed ambiguous results for the FA model.

**Figure 3:**
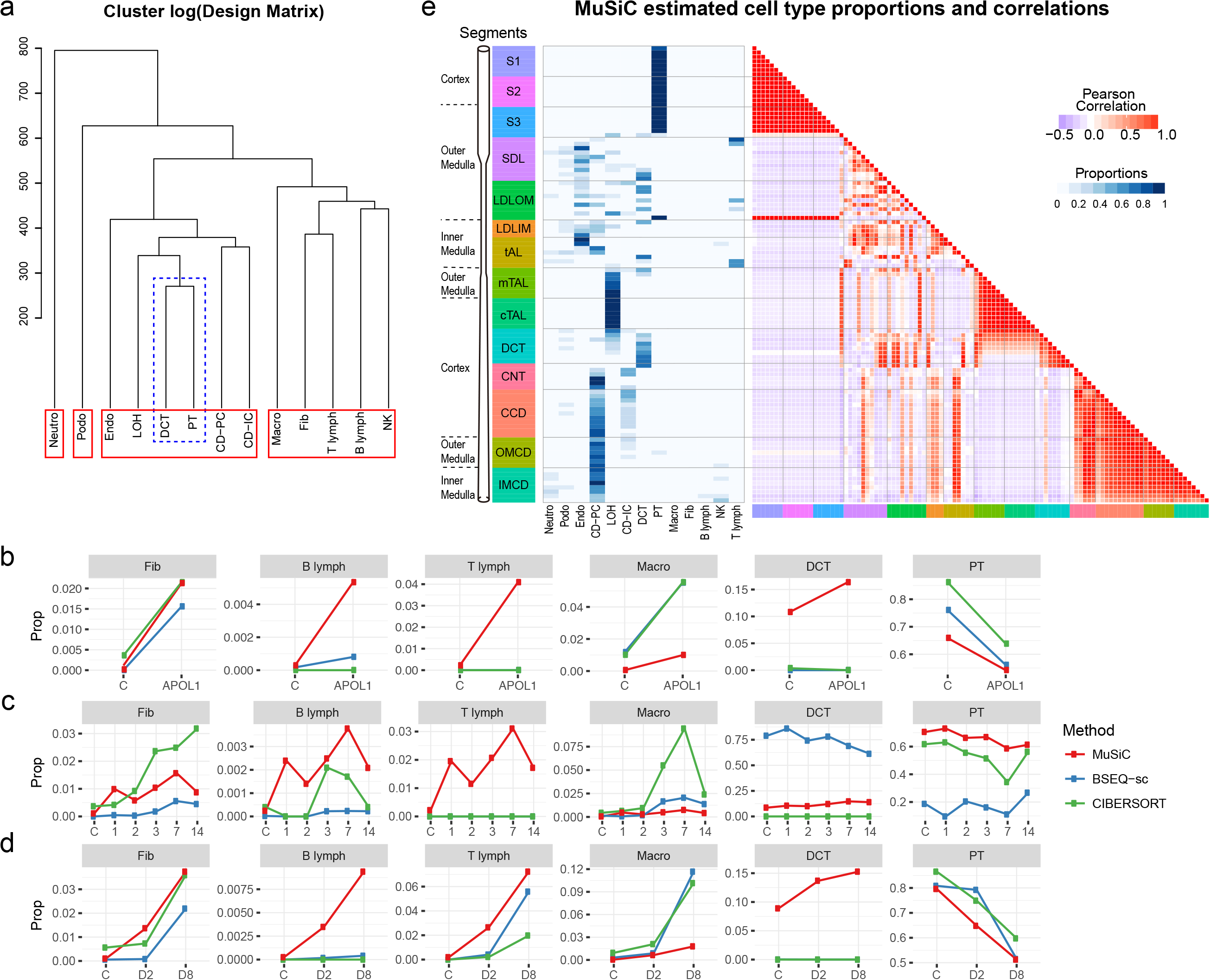
Cell type composition in kidney of mouse CKD models and rat. **a**. Cluster dendrogram showing similarity between 13 cell types that were confidently characterized in Park et al. Abbreviations: Neutro: neutrophils, Podo: podocytes, Endo: endothelials, LOH: loop of Henle, DCT: distal convolved tubule, PT: proximal tubule, CD-PT: collecting duct principal cell, CD-IC: CD intercalated cell, Macro: macrophages, Fib: fibroblasts, NK: natural killers. **b, c and d**. Average estimated proportions for 6 cell types in bulk RNA-seq samples taken from 3 different studies, each study based on a different mouse model for chronic kidney disease. Results from three different deconvolution methods (MuSiC, BSEQ-sc and CIBERSORT) are shown by different colors. **Supplementary Figure 5a-c** show complete estimation results of all 13 cell types. **b**. Bulk samples are from Beckerman et al., who sequenced 6 control and 4 APOL1 mice. **c**. Bulk data are from Craciun et al.^9^, where samples are taken before (C) and at 1, 2, 3, 7, 14 days after administering folic acid. Line plot shows cell type proportion changes over time (days), averaged over 3 replicates at each time point. **d**. Bulk data are from Arvaniti et al.^10^, where samples are taken from mice after Sham operation (C), 2 days after UUO operation (D2), and 8 days after UUO operation (D8). The average proportions at each time point are plotted. **e**. MuSiC estimated cell type proportions of rat renal tubule segments. The estimated cell type proportions (left) and the proportions correlations between samples (right) are shown as heatmap. Segment names are color coded and aligned according to their physical positions along the renal tubule. **Supplementary Figure 6a-c** show NNLS, BSEQ-sc and CIBERSORT results. Segment name abbreviation: S1: S1 proximal tubule; S2: S2 proximal tubule; S3: S3 proximal tubule; SDL: Short descending limb; LDLOM: Long descending limb, outer medulla; LDLIM: Long descending limb, inner medulla; tAL: Thin ascending limb; mTAL: Medullary thick ascending limb; cTAL: Cortical thick ascending limb; DCT: Distal convoluted tubule; CNT: connecting tubule; CCD: Cortical collecting duct; OMCD: Outer medullary collecting duct; IMCD: Inner medullar collecting duct.

Distal convolved tubule cells (DCT) are known to be the second most numerous cell type in kidney, with an expected proportion of ~10-20%^1^. Yet, CIBERSORT did not detect DCT in any of the three bulk datasets; BSEQ-sc missed it in two datasets and grossly over-estimated its proportion in the third dataset at the cost of a grossly underestimated PT proportion. This is due to the high similarity between DCT and PT, observable in **Figure 3a**. Through its tree-guided recursive algorithm, MuSiC first estimates the combined proportion of kidney cell types versus immune cell types using stable genes for these two large groups, and then zooms in and deconvolves the kidney cell types using genes re-selected for each kidney cell type. This allows MuSiC to successfully separate PT and DCT cells in all three bulk datasets, recovering a consistent DCT proportion between 8-20%, matching expectations. Interestingly, unlike for PT, the proportion of DCT cells show a consistent increase with disease progression across all three mouse models. This may seem counterintuitive given that loss of kidney function is expected to be associated with the loss of kidney cell types. But given the substantial drop of the dominant PT cell type, the proportion of DCT cells relative to the whole may increase, even if its absolute count drops.

Next, consider immune cells, known to play a central role in the pathogenesis of CKD. MuSiC found the largest immune sub-type to be macrophage, and all methods detected the expected increase of macrophage proportion with disease progression. Apart from this, MuSiC also found fibroblasts, B-, and T-lymphocytes to increase in proportion with disease progression, giving a consistent immune signature that is reproduced across mouse models. These findings are consistent with clinical and histological observations, indicating tissue inflammation is a consistent feature of kidney fibrosis. Such reproducible signatures were not found by other methods, which show much less agreement across mouse models.

Finally, to illustrate MuSiC’s cross-species applicability, we used the mouse kidney scRNA-seq reference from Park et al.^1^ to deconvolve the bulk rat RNA-seq data from Lee et al.^13^, which contains 105 samples obtained from 14 segments spaced along the renal tubule. We mapped samples to their physical locations, and computed correlations between their cell type proportions (**Figure 3e**). Reassuringly, cell types recovered by MuSiC for each segment agree with knowledge about the dominant cell type at its mapped position, e.g. DCT cells come from the DCT region. Correlation between samples is also high within anatomically distinct segments.

Knowledge of cell type composition in disease relevant tissues is an important step towards the identification of cellular targets in disease. Although most scRNA-seq data do not reflect true cell type proportions in intact tissues, they do provide valuable information on cell type-specific gene expression. Harnessing multi-subject scRNA-seq reference data, MuSiC reliably estimates cell type proportions from bulk RNA-seq. As bulk tissue data are more easily accessible than scRNA-seq, MuSiC allows the utilization of the vast amounts of disease relevant bulk tissue RNA-seq data for elucidating cell type contributions in disease.

## Acknowledgments

This work was supported by the following funding: NIH R01HG006137 (to N.R.Z.); NIH R01GM125301 (to N.R.Z., M.L.); NIH R01GM108600 (to M.L.); NIH R01HL113147 (to M.L.); NIH R01DK076077 (to K.S., M.L.); NIH R01DK087635 (to K.S.); NIH R01DK105821 (to K.S.); ADA postdoctoral fellowship (to J.P.).

## Author Contributions

This study was conceived of and led by N.R.Z. and M.L. Jointly with N.R.Z. and M.L., X.W. designed the model and estimation algorithm, implemented the MuSiC software, designed the *in silico* experiments, and led the data analysis. J.P. and K.S. performed the mouse scRNA-seq experiment and provided scientific insight on chronic kidney disease and data interpretation. X.W., N.R.Z. and M.L. wrote the paper with feedback from J.P. and K.S.

## Online Methods

### MuSiC model set-up

In this section, we derive the relationship between gene expression in bulk tissue and cell type-specific gene expression in single cells. This relationship forms the basis of our regression-based deconvolution. For gene *g*, let *X*_*jg*_ be the total number of mRNA molecules in subject *j* of the given tissue, which is composed of *K* cell types.

Then, 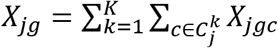, where *X*_*jgc*_ is the number of mRNA molecules of gene *g* in cell *c* of subject *j*, and 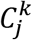 is the set of cell index for cell type *k* in subject *j* with 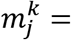 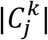 being the total number of cells in this set. The relative abundance of gene *g* in subject *j* for cell type *k* is

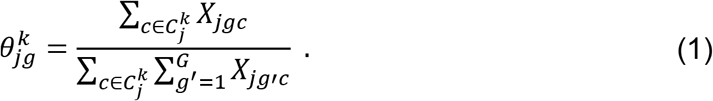

We can show that

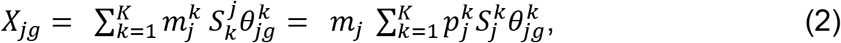

where, for subject *j*, 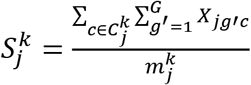 is the average number of total mRNA molecules for cells of cell type *k* (also referred to as “cell size” below), 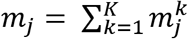 is the total number of cells in the bulk tissue, and 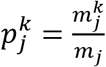 is the proportion of cells from cell type *k*. Let 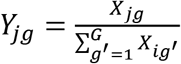 be the relative abundance of gene *g* in the bulk tissue of subject *j*. Equation (2) impiles

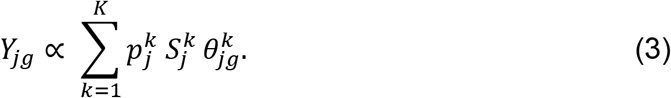

Thus, across *G* genes in subject *j*, we have

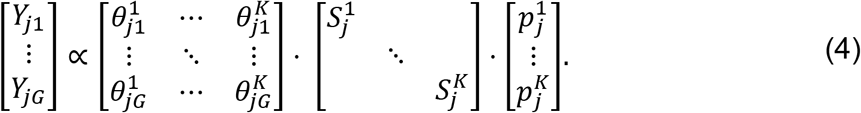

The goal of MuSiC is to estimate 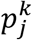 using data from scRNA-seq and bulk RNA-seq.

### Model assumptions

If scRNA-seq data were available for subject *j*, we would be able to obtain the cell size factor 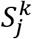 and cell type-specific relative abundance 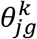. With bulk RNA-seq data in subject *j*, we get the bulk tissue relative abundance *Y*_*jg*_, and, if 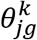 and 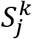 were known, we would be able to perform a regression to estimate 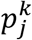. However, since scRNA-seq is still costly, most studies cannot afford the sequencing of a large number of individuals using scRNA-seq. To make deconvolution possible for a broader range of studies, it is desirable to utilize cell type-specific gene expression from other studies or from a smaller set of individuals in the same study. This is feasible under the following two assumptions: (A1) Individuals with scRNA-seq and bulk RNA-seq are from the same population, with their cell-type specific relative abundances 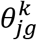 in equation (1) following the same distribution with means 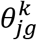 and variances of 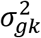,

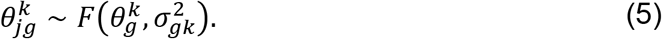

Under this assumption, deconvolution can use available single cell data from other subjects or even subjects from other studies as reference for cell type proportion estimation. (A2) The ratio of average cell size 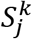 across cell types are the same regardless of subjects and studies

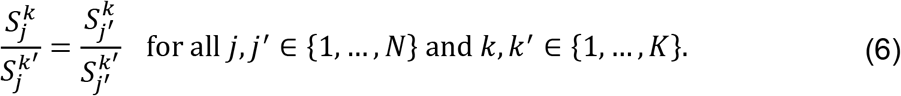

The second assumption allows us to replace 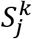 by a common value *S*^*K*^ across subjects. In MuSiC, we use the average cell size and relative abundance across all subjects from the scRNA-seq data to estimate 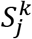 and 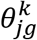.

### Cell type proportion estimation

To estimate cell type proportions 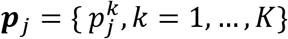 we need to consider two constraints: (C1) Non-negativity: 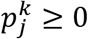 for all *j,k*; (C2) Sum-to-one: 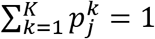 for all *j*. Because the bulk tissue and single-cell relationship derived in equation (5) is a “proportional to” relationship, to satisfy the (C2) constraint, we need a normalizing constant *C* so that

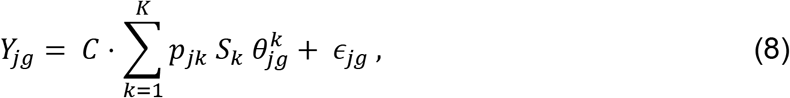

where 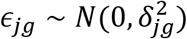 represents bulk tissue RNA-seq gene expression measurement noise. When cell type proportions 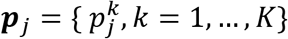 and subject-specific relative abundances 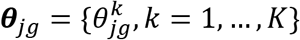 are known, the variance of bulk tissue gene expression measurement is

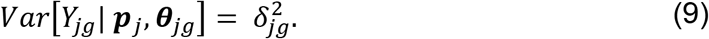

Given only cell type proportions, the variance is

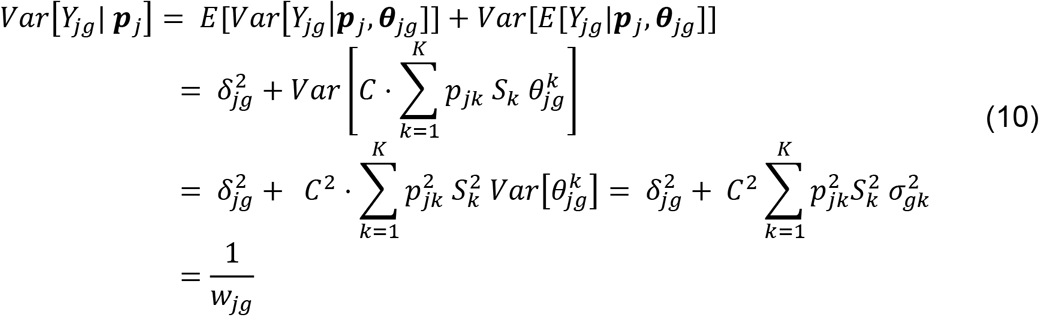

Because of the heteroscedasticity of gene expression over genes, including the weight *w*_*jg*_ can improve estimates. Since 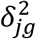 is unknown, we will estimate *w*_*jg*_ iteratively, initialized by NNLS.

MuSiC is a weighted non-negative least squares regression (W-NNLS), which does not require pre-selected marker genes. Indeed, the iterative estimation procedure automatically imposes more weight on informative genes and less weight on non-informative genes. Because it is a linear regression-based method, genes showing less cross cell type variations will have low leverage, thus having less influence on the regression, whereas the most influential genes are those with high weight and high leverage. To illustrate this point, we also performed benchmarking experiments to show that applying MuSiC using all genes gives more accurate results than applying MuSiC using pre-selected marker genes, thus demonstrating that MuSiC’s weighting scheme makes marker gene pre-selection unnecessary (**Supplementary Figure 1c, Supplementary Figure 2**).

### Recursive tree-guided deconvolution for closely related cell types

Complex solid tissues often include closely related cell types with similar gene expression levels. Correlation in gene expression can lead to collinearity, making it difficult to reliably estimate cell type proportions, especially for less frequent and rare cell types. Although the collinearity problem can be improved by selecting marker genes through support vector regression, as is done in CIBERSORT^3^ and BSEQ-sc^4^, these approaches still have limited power to resolve similar cell types. In MuSiC, we introduce a recursive tree-guided deconvolution procedure based on a cell type similarity tree, which can be easily obtained through hierarchical clustering. In stage 1 of this procedure, cell types in the design matrix are divided into high-level clusters by hierarchical clustering with closely related cell types clustered together. Proportion for these cell type clusters are estimated using genes with small intra-cluster variance (cluster-stable genes) using the above described W-NNLS. In stage 2, for cell types in each cluster, the cell type proportions are estimated using W-NNLS with genes displaying small intra-cell type variance, subject to the constraint on the pre-estimated cluster proportions. If necessary, more than 2 stages of recursion can be applied, with each stage separating the cell types within each large cluster into finer clusters, and using cluster-stable genes to do W-NNLS subject to the constraint that fixes higher-level cluster proportions.

To illustrate this recursive tree-guided deconvolution procedure, we start with a simple case with four cell types and *G* genes. Let *X*_1_,*X*_2_,*X*_3_,*X*_4_ represent cell type-specific expression in the design matrix, obtained from scRNA-seq, and let *Y* be the gene expression vector in the bulk RNA-seq data. The relationship of bulk and single-cell data can be written as

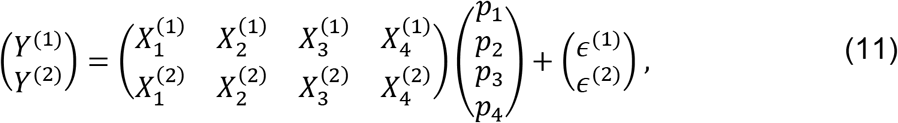

where the superscripts (1) and (2) indicate two sets of genes. Suppose the four cell types are grouped into two clusters, (*X*_1_,*X*_2_) and (*X*_3_,*X*_4_). The first set of genes are those showing small intra-cluster variance in gene expression, that is, 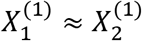 and 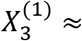 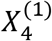, whereas the second set of genes are the remaining genes.

<u>*Stage 1*</u>: Estimate cluster proportions *π*_1_ = *p*_1_ + *p*_2_ and *π*_2_ = *p*_3_ + *p*_4_,

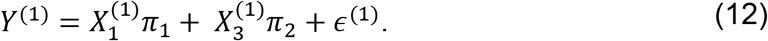 The cluster proportions, 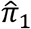 and 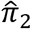 are estimated by W-NNLS using intra-cluster homogenous genes.
<u>*Stage 2*</u>: Estimate cell type proportions (*p*_1_, *p*_2_, *p*_3_, *p*_4_),

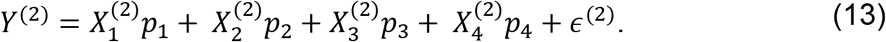 The cell type proportions are estimated by W-NNLS using the remaining genes subject to the constraint that

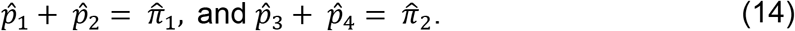

### Construction of benchmark datasets and evaluation metrics

To evaluate MuSiC and compare with other deconvolution methods, we need bulk RNA-seq data with known cell type proportions. Therefore, we construct artificial bulk tissue data from a scRNA-seq dataset in which the bulk data is obtained by summing up gene counts from all cells in the same subject. Relative abundance is calculated by equation (1). The true cell type proportions in the artificial bulk data can be directly obtained from the scRNA-seq data and this allows us to use this artificially constructed bulk data as a benchmark dataset to evaluate the performance of different deconvolution methods. Denote the true cell type proportions by ***p*** and the estimated proportions by 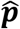. Deconvolution methods are evaluated by the following metrics.

i. Pearson correlation, 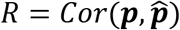.
ii. Root mean squared deviation, 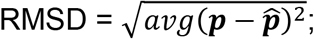
iii. Mean absolute deviation, 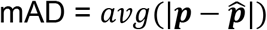.

